# Structural investigation of Mycobacterial protein complexes involved in the stationary phase stress response

**DOI:** 10.1101/2020.03.04.976621

**Authors:** Angela M Kirykowicz, Jeremy D Woodward

## Abstract

Large protein complexes play key roles in mediating biological processes in the cell. Little structural information is known on the protein complex mediators governing persistence in the host for *Mycobacterium tuberculosis* (*Mtb*). We applied the ‘shotgun EM’ method for the structural characterisation of protein complexes produced after exposure to stationary phase stress for the model Mycobacterium, *M smegmatis* (*Msm*). We identified glutamine synthetase I, essential for *Mtb* virulence, in addition to bacterioferritin, critical for *Mtb* iron regulation, and encapsulin, which produces a cage-like structure to enclose target proteins. Further investigation found that encapsulin carries dye-decolourising peroxidase (DyP), a potent protein antioxidant, as the primary cargo during stationary phase stress. Our ‘proof-of-concept’ application of this method offers insight into identifying potential key-mediators in *Mtb* persistence.

## Introduction

Protein machinery allows for the vast biochemical capabilities of cells, driving replication, repair, and response to environmental stresses. The pathogenic bacteria, *Mycobacterium tuberculosis* (*Mtb*), relies on a range of critical protein machinery to evade and utilise the host immune response (1). Although some studies have shown potential key mediators in *Mtb* survival in the host (2–4) there is a lack of structural information to guide drug discovery or elucidate biological function.

Structural data provides compelling evidence for the existence of protein complexes which are physiologically relevant to the cell. In addition, this data offers valuable information on subunit composition and mechanism of interaction, which yields biological insight governing its potential actions (5). Single-particle transmission electron microscopy (TEM) is a powerful method to reconstruct large protein complexes. The method has been successfully used to solve the structures of protein complexes in a range of organisms from homogenous (6) as well as heterogenous (7–10) samples. Combining 3D reconstruction of protein complexes from negative stain EM images with information obtained from mass spectrometry data (‘shotgun EM’) (10) allows for a potentially high-throughput approach to finding complexes which could play a key role in *Mtb* pathogenesis.

Although the ‘shotgun EM’ approach offers a faster method for solving the structures of a range of protein complexes without the need for extensive purification (11), the approach still needs to be tested and adapted for the organism of application (9,10,12). Here, we present our ‘shotgun EM’ methodology for the purification and TEM 3D reconstruction of Mycobacterial protein complexes from the model organism, *M smegmatis* (*Msm*), after exposure to stationary phase stress which is known to induce a protective effect against subsequent exposure to oxidative stress (13). We identified three protein complexes (glutamine synthetase I (GSI) (E.C 6.3.1.2), bacterioferritin (BrfA) (E.C 1.16.3.1), and encapsulin); the initial identity assigned by mass spectrometry, as well as the 3D model and subunit composition, was validated by fitting of homologous crystallographic structures into the low-resolution density. We demonstrate that encapsulin encloses dye-decolourising type peroxidase (DyP) (E.C 1.11.17), a potent enzymatic anti-oxidant (14), as the main cargo during stationary phase stress. Furthermore, analysis of our encapsulated DyP shows that it binds on the encapsulin three-fold axis, validating the relationship between cargo binding and substrate access *in vivo*.

## Results

### Partial fractionation in combination with *in silico* purification yields three-dimensional structures

We induced production of protein complexes potentially involved in the oxidative stress response by growing *Msm* culture to the end of stationary phase (13). As an initial partial fractionation step, we applied ammonium sulfate precipitation to cell lysate. We noted that the EM micrographs displayed a large amount of soluble aggregated protein. Thus, we opted to use a manual picking strategy to limit the amount of “junk” particles in the dataset and improve classification of true (non-aggregated) protein complexes. However, the resolution of separation was too low to build models after applying *in silico* purification by particle picking and classification (**SI Figure 1**). We thus opted for either partial fractionation by anion exchange or sucrose-cushioning. We applied a size cut-off (>100 kDa) when concentrating obtained peaks for EM analysis. Manually picked particles were used to generate separate EM models using the symmetry information gathered from the average in the particle stack (**Figure 1**).

**Figure 1.**
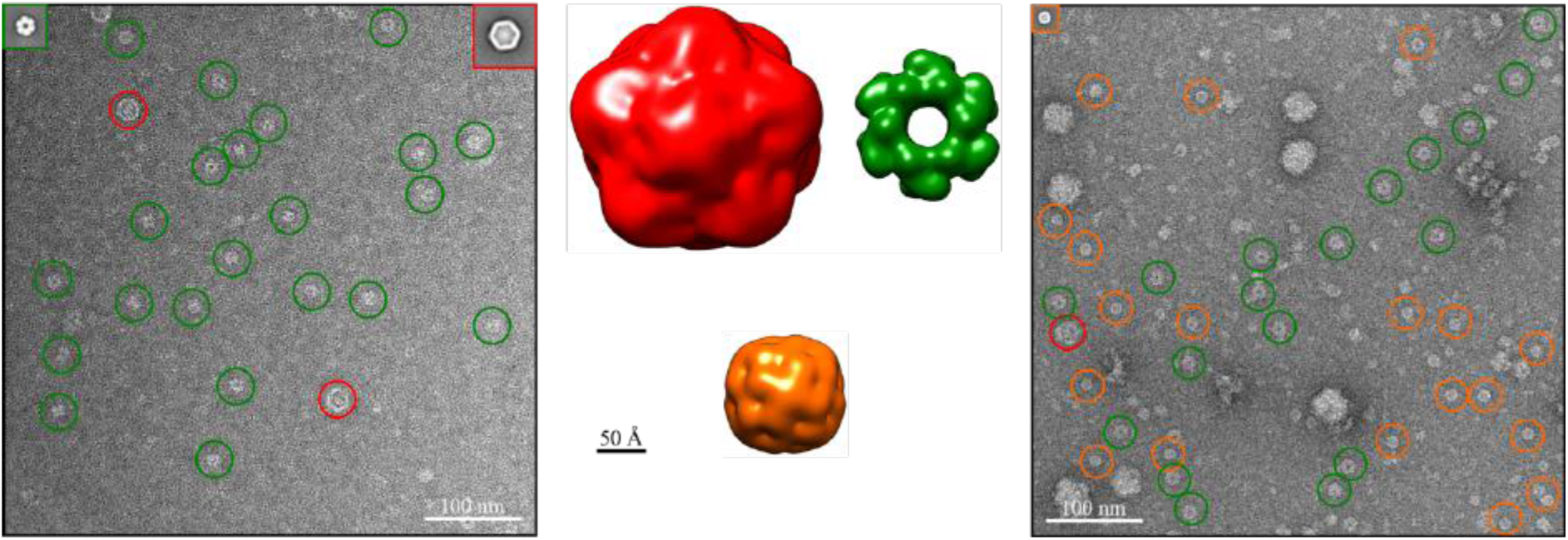
Partial fractionation and *in silico* purification. Three protein complexes were partially purified from anion exchange chromatography (left) or sucrose cushioning (right). Particles were further purified *in silico* through particle picking and classification. This lead to reconstructions for encapsulin (cfp29) (red), glutamine synthetase I (GSI) (green), and bacterioferritin (BrfA) (orange). Particles for encapsulin and GSI were also found in sucrose density purification, but the particles were of too poor quality for averaging and further reconstruction.

### Identification by mass spectrometry and verification by model fitting

We succeeded in unambiguous identification of our models through a combination of information from mass spectrometry and fitting of homologous structures into our low-resolution models (~25 Å) (**Figure 2**). We first used the EM models to estimate molecular weight, which was roughly size matched to bands in a native PAGE gel of anion exchange fractions. Analysis by liquid chromatography tandem mass spectrometry (LC–MS/MS) showed that peptides for glutamine synthetase I (GSI) and 29 kDa antigen Cfp29 (encapsulin) showed respective highest number of unique peptides and coverage (**SI Table 1**).

**Figure 2.**
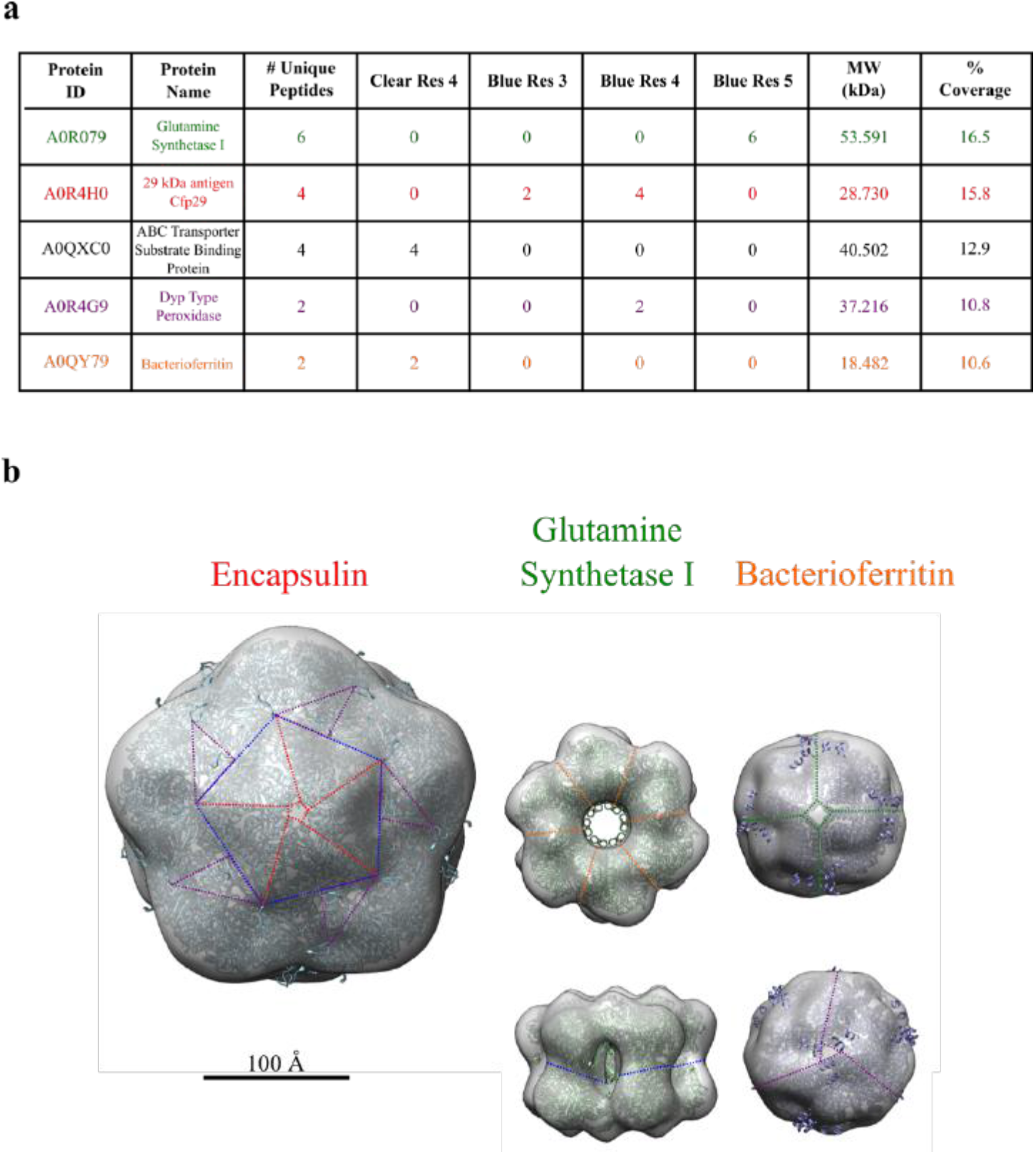
Mass spectrometry and model fitting. (**a**) We analysed bands cut out from clear and blue native gels of solubilised membrane-bound protein and analysed them by LC-MS/MS. Single MS/MS peptide hits, or those which were likely to be degraded, or present in blank runs were manually removed from analysis. For full data see SI Table 2. (**b**) We searched for crystal structures with the same symmetries in the PDB. Encapsulin from *Thermotoga maritima* (3dkt) (19), and glutamine synthetase I (1hto) (18) and bacterioferritin B (3uno) (20) from *Mycobacterium tuberculosis* were found and docked into the appropriate density. Crystal structures have good correspondence to the density and symmetry axis on each low-resolution structure. The fit was evaluated by cross-correlation. The 6-fold (orange), 4-fold (green), 5-fold (red), 3-fold (purple), and 2-fold (blue) symmetry axes are shown as appropriate for each structure.

We noted extra density present in the encapsulin particles; furthermore, it is known that recombinant *Mtb* encapsulin can contain three types of cargo protein: dye-decolourising type peroxidase (DyP), bacterioferritin B (BrfB), or 7,8-dihydroneopterin aldolase (FolB) (15). However, we found no plausible hits which could identify the extra density in the LC–MS/MS data (**SI Table 1**). We looked at both GSI and encapsulin in the Mycobrowser (https://mycobrowser.epfl.ch/) database (16) and found that they are both membrane associated. Hence to find a potential cargo lead for encapsulin, we isolated the membrane fraction of *Msm,* and ran the resolubilised material on either blue or clear native PAGE; to not introduce bias, all bands were cut out and analysed by LC–MS/MS (**Figure 2a; SI Table 2**). Both GSI and encapsulin (Cfp29) were the major peptides found in different blue native bands (**Figure 2a**). Bacterioferritin (BrfA) was also found as a minor peptide in two clear native bands (**SI Table 2**). The only major peptide which was co-found exclusively with Cfp29 was DyP **(Figure 2a; SI Table 2**). Thus, it seemed likely that this was the potential cargo protein for *Msm* encapsulin. To confirm the result, we ran SDS-PAGE on gel filtration fractions confirmed by EM analysis of micrographs to harbour encapsulin particles and cargo (**SI Figure 2**). We cut out a band which corresponded to the approximate size of DyP (~40 kDa) and analysed it by mass spectrometry. Hits which did not match the mass of the band calculated using a standard calibration curve for the gel were excluded from analysis. DyP was found with 6.4% coverage **(SI Table 3**). None of the other known cargo proteins were found.

To verify protein identities assigned by LC–MS/MS, we searched the Protein Data Bank (PDB) (17) for available solved structures which matched the symmetry of our low-resolution models. We fitted hits manually at first, and optimised by local fitting refinement and checked by crosscorrelation; GSI (1hto) (0.8795) (18), encapsulin (3dkt) (0.7319) (19), and bacterioferritin (3uno) (0.9361) (20). Fitted crystallographic structures also showed good correspondence to the symmetry of our models (**Figure 2b**), validating the subunit composition.

### Binding of DyP cargo on encapsulin three-fold axis and possible export

It is known that encapsulins function to enclose target proteins via a unique C-terminal extension on the encapsulated protein (15, 21). Additionally, both *Msm* DyP and BrfB, but not *Msm* FolB or BrfA, contain C-terminal extensions (15, 22) (**SI Figure 3**). However, only DyP was identified in our previous LC–MS/MS results. Inspection of *Msm* encapsulin particles showed that BrfB could potentially be present (**Figure 3a**).

**Figure 3.**
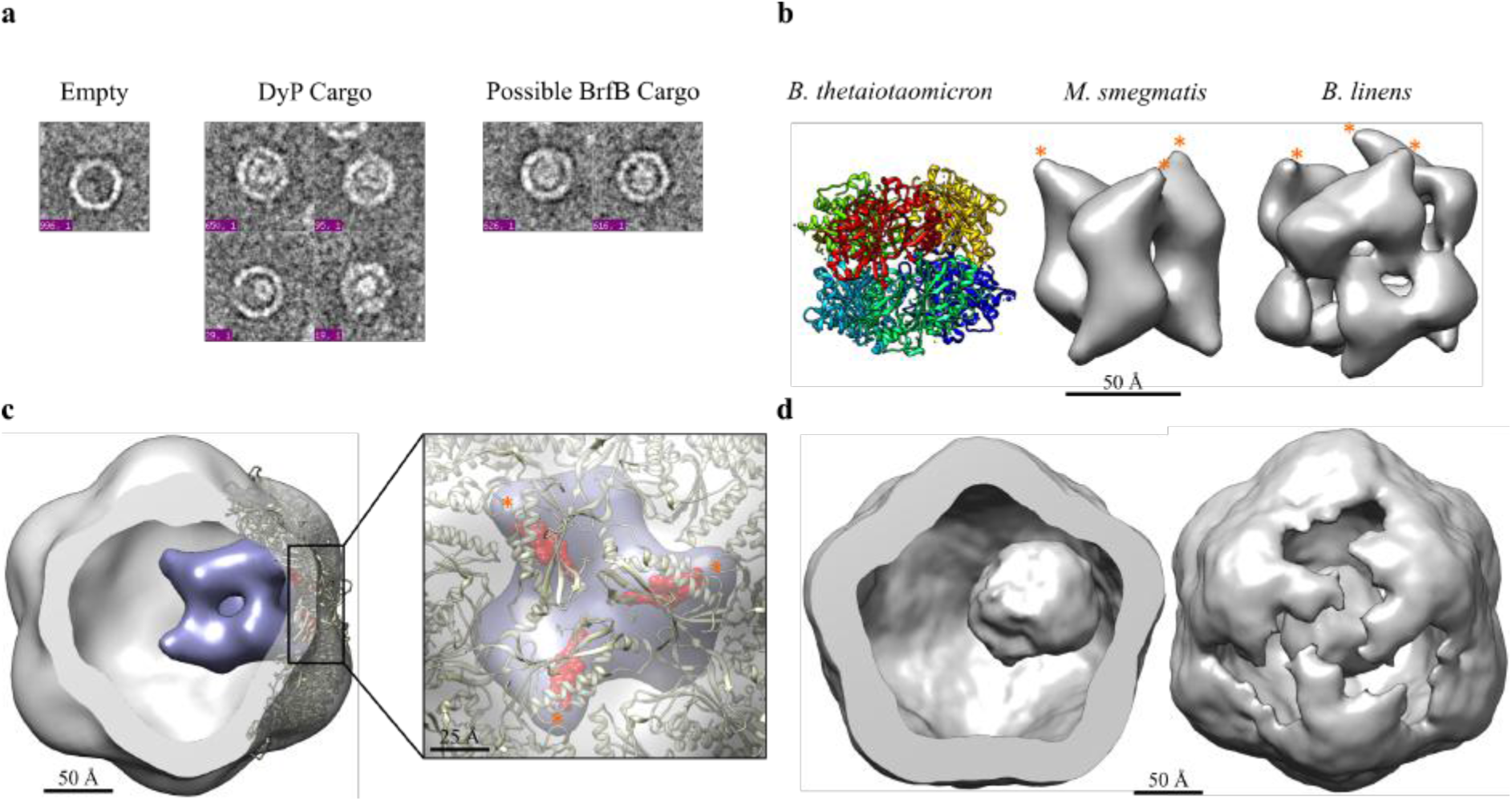
Identification of the primary cargo of *M smegmatis* enapsulin in response to stationary phase stress. **(a)** Encapsulin particles can either be empty or contain either DyP-type peroxidase or bacterioferritin B cargo. Bacterioferritin B appears square-like, owing to its octahedral symmetry, while the 3- and 2-fold symmetries of DyP can be seen depending on the particle orientation. **(b)** Prelimary reconstruction of encapsulated DyP (middle) compared to other solved DyP crystal (left, pdb code 2gvk) (46) or low-resolution EM (right, emd-1530) (47) structures. The position of the C-terminal extension is starred (orange). Note that the C-terminal extension for *B. thetaiotaomicron* DyP was not built into the crystal structure. **(c)** Model of DyP (purple) docked into the 3-fold axis of encapsulin (grey) fills a substantial part of the lumen. It is clear from the docking that only a single DyP hexamer is likely to be accommodated. The C-terminal extension (red) from the *T. maritima* encapsulin crystal structure (brown) (pdb code 3dkt) (19) aligns well with the location of the missing DyP extension (starred, orange). **(d)** Application of three-fold symmetry reveals DyP on the three-fold symmetry axis (left), corresponding to the expected position based on the docking model in (c). The density of encapsulin has been partially stripped (right) to better show DyP on the three-fold axis.

Since all captured proteins are membrane associated, and there is previous indication that *Mtb* Cfp29 (encapsulin) can be exported (23), we partially purified cell-culture filtrate by anion exchange. We detected the presence of encapsulin and cargo by EM micrographs, but failed to detect an SDS-PAGE band corresponding to the expected size of BrfB (**SI Figure 4**). Thus, it seemed unlikely that BrfB is the main encapsulin cargo. Furthermore, it has been proposed that any bound cargo lies at a specific symmetry axis on encapsulin, corresponding to the symmetry of the bound protein (e.g two-fold, three-fold, or five-fold axis) (21) (**SI Figure 3**). DyP could potentially bind on the encapsulin three-fold axis (**SI Figure 3**). To address this question, we manually removed empty/deformed particles or those which may contain BrfB, and reconstructed the encapsulated cargo by masking out the shell. The obtained model is highly consistent with previously solved peroxidase structures, showing a hexameric subunit arrangement, with the C-terminal extension lying on the three-fold axis (**Figure 3b**) (14,21). By docking our model back into *Msm* encapsulin density, we found that the three-fold axis showed good alignment with the tethering C-terminal extension found in *T. maritima* encapsulin crystal structure; only one hexamer can be accommodated in the lumen (**Figure 3c**). To confirm the possibility of DyP binding at a specific symmetry axis, we applied C3 symmetry to unmasked 2D projections; this revealed density clustered to the encapsulin three-fold axis, which is likely to be DyP (**Figure 3d**). Thus, DyP was confirmed to be the primary cargo *in vivo* of *Msm* encapsulin under stationary phase stress and there is some evidence that it binds specifically to the three-fold axis, a result which has also been found for *B. linens* encapsulin (24).

## Discussion

We obtained three protein complexes (GSI, bacterioferritin, and encapsulin) by ‘shotgun EM’ after *Msm* was exposed to stationary phase stress. Identification of these initially unknown protein complexes proved to be particularly challenging and relied on a combination of LC– MS/MS of native PAGE gel bands and fitting of homologues crystal structures. Our LC– MS/MS results were confirmed by multiple partial fractionation strategies. However, since it is known that peptide abundance does not strictly correlate to protein abundance (25), and because the number of unique identified peptides for all three proteins was low, we opted to verify the mass spectrometry results by finding homologous crystal structures in the PDB to fit into our low-resolution models. The crystal structure homologues matched the mass spectrometry data, lending confidence to the result. One drawback of this approach is that it relies on both the availability of a homologue in the PDB and the conservation of the quaternary structure. Use of higher resolution structures could potentially eliminate the need for an existing PDB structure to verify the mass spectrometry results.

Under stationary phase stress, *Msm* encapsulin appears to primarily enclose DyP, a potent protein antioxidant (14). The main role of BrfB, another potential encapsulated protein, is to relieve iron toxicity (26). EM models provide a good indication of the relative abundance of binding partners in a protein complex; more abundant members will dominate the density (7,27). We noted that the majority of particles appeared to contain DyP rather than BrfB cargo; our reconstruction showed a typical peroxidase structure, which was unlikely to have been biased by any remaining BrfB particles. The production of encapsulated DyP may have functional significance in *Msm*, as one of the mechanisms in which *Msm* has increased resistance to oxidative stress after growth in stationary phase (13). Furthermore, we found that *in vivo* DyP is bound on the encapsulin three-fold axis, implying that a specific pore is involved in the entry of the hydrogen peroxide substrate. Since it is also possible for BrfB to be bound at the three-fold axis, and reaction of hydrogen peroxide with iron can produce harmful hydroxyl radicals via the Fention reaction (28), it remains to be further investigated how encapsulin regulates substrate entry.

GSI is known to be essential in *Mtb* virulence (29) and growth in macrophages (30), while bacterioferritin (BrfA) is crucial for iron regulation (31). We confirmed that all three proteins are membrane-associated by isolating the membrane fraction and analysing solubilised proteins by LC–MS/MS. This led us to examine the cell-culture filtrate and we found some evidence for the presence of encapsulin and DyP cargo. This leads to an interesting possibility that these proteins could potentially be exported during stationary phase stress, as suggested by previous investigators (23,32). Thus, we have isolated three protein complexes by the new ‘shotgun EM’ methodology which can all be implicated in *Msm* response to stationary phase stress, and also play important roles in *Mtb* persistence in the host. Our proof-of-concept study has the potential to be adapted to find more protein complexes involved in *Mtb* pathogenesis.

## Experimental Procedures

### Bacterial Growth

Culture (Middlebrook 7H9 media supplemented with 0.2% glucose, 0.2% glycerol, and 0.05% Tween-80) of *Msm* groELΔC (33) was inoculated with 1/100 starter culture and grown at 37°C with shaking (120 rpm) to the end of stationary phase (~4–5 days). Cells were harvested through centrifugation (4000g for 30 minutes) at 4°C. The pellet was stored at −80°C.

### Cell Lysis and Ammonium Sulphate Precipitation

The pellet was thawed and resuspended in 25 mL of lysis buffer (50 mM Tris-HCl, 300 mM NaCl, pH 7.2) with protease inhibitor cocktail. Cells were lysed through 4 x (15 seconds on, 15 seconds off for 4 minutes) on ice. Cell debris was pelleted by centrifugation (20,000g for 1 hour) at 4°C.

Ammonium sulphate cuts were completed on the filtered (0.45 μm) supernatant (<40%, 40-50%, 50-60%, and >60%). For each cut, the ammonium sulphate was added slowly on ice with continual stirring and incubated for 30 minutes before centrifuging at 9000g for 15 minutes. Pellets were clarified by re-suspending in 20 mL of gel filtration buffer (50 mM Tris-HCl, 200 mM NaCl, pH 8.0) and centrifuged at 20,000g for 10 minutes at 4°C. The ammonium sulphate cuts were then buffer exchanged to gel filtration buffer using an Amicon^®^ spin-filter with a 100 kDa cut-off (Merck, Darmstadt, Germany).

### Anion Exchange

20 mL HiPrep Q FF 16/10 column (GE Healthcare Life Sciences, Massachusetts, USA) was equilibrated with 5–10 column volumes of start buffer (20 mM Tris-HCl, 20 mM NaCl, pH 8.0). Samples were then eluted with 0.5 M NaCl (3 CV), before applying a gradient of 0.5 – 1 M NaCl (19.5 CV). The flow rate was 5 mL/min with 60 fractions collected. Fractions were stored at 4°C.

### Gel Filtration

The PWXL5000 column (Tosoh Biosciences, Tokyo, Japan) was equilibrated with gel filtration buffer and run at a flow rate of 0.5 mL/min for 1 column volume. Fractions were stored at 4°C.

### Sucrose Cushioning

The method was adapted from Peyret (2015) (34) with the following modifications: a double sucrose cushion consisting of 25% (top layer) and 70% (bottom layer) sucrose made in sodium phosphate buffer (pH 7.4). The sample was spun at 170,462g for 5 hours. The layer just above the 70% cushion was extracted and buffer exchanged to gel filtration buffer using an Amicon^®^ spin-filter with a 100 kDa cut-off (Merck, Darmstadt, Germany).

### Membrane Preparation and Electrophoresis

2 L of *Msm* culture was grown as described and membranes prepared for electrophoresis as described previously (35,36). For clear native PAGE, standard continuous Tris-Glycine (pH 8.8) system was used.

### Negative Stain Electron Microscopy

Samples were pipetted onto a glow-discharged (in air) copper grid and washed/stained with 5 rounds of 2% uranyl acetate before being air-dried. Images were taken using the Tecnai F20 transmission electron microscope (Phillips/FEI, Eindhoven, The Netherlands) fitted with a CCD camera (4k x 4k) (GATAN US4000 Ultrascan, USA) at 200 kV under normal dose conditions with a defocus of 2.00 μm at the appropriate magnification. The sampling rate was 2.11 or 3.84 Å/pixel.

### Class Averages

Class averages were produced in Appion (37) from manually picked particles. Briefly, the Contrast Transfer Function (CTF) was estimated using ACE2 (38) and poor images excluded based on the presence of astigmatism, bad staining, or noticeable microscope drift. A stack was created with CTF correction (ACE2 Phaseflip of whole image) (38). A Spider reference-free alignment (39) was completed, averaging all particles in the stack. Afterwards, Spider Coran classification (39) was completed using appropriate settings. Either K-means or hierarchical clustering was completed using selected eigen images.

### Reconstruction

All reconstructions were completed in the Appion pipeline (37). Particles were binned by a factor of 2 for a sampling of 2.11 Å/pixel. The appropriate number of classes was used to complete an initial reconstruction using EMAN Common Lines (40) with the appropriate symmetry imposed. The model was then refined using EMAN model refinement (40) for 26 iterations with the appropriate symmetry imposed; 20 iterations was used for GSI. Angular sampling was as follows: 5 iterations of 10°, 5 iterations of 8°, 10 iterations of 5°, and 6 iterations of 3°. For GSI, the angular sampling was 20 iterations of 5°.

### Reconstruction of Encapsulated Dye-Decolourising Peroxidase

A sub-stack was created in which particles that were broken, deformed, or may contain BrfB were deleted. This left 207 particles. Encapsulin was masked out using a rectangular box with a Gaussian drop-off in intensity. Class averages were produced as described previously using hierarchical clustering. An initial model was created using EMAN Common Lines (40) with C3 symmetry imposed. This model was refined using EMAN projection-matching (40) with D3 symmetry imposed for 26 iterations. For refinement, a 15 Å low pass filter was used and a mask radius of 70 Å applied. Angular sampling rate was used as described previously.

### Mass Spectrometry

Samples were sent for MS either to the Blackburn Group (in-gel native PAGE LC-MS/MS) (University of Cape Town, South Africa) or to the Yale MS & Proteomics Resource (in gel SDS-PAGE LC-MS/MS) (Yale School of Medicine, New Haven, USA). Samples were digested with trypsin and analysed on an LTQ Orbitrap (ThermoScientific, Massachusetts, USA). MS/MS spectra were searched using the Mascot algorithm (41). Peaks with a charge state of +2 or +3 were located first using a signal-to-noise ratio of >1.2. Potential peaks were screened against the NCBInr or SWISS-PROT (42) databases.

### Bioinformatics

Obtained EM models were imported into UCSF-Chimera (43) and set to the correct voxel size. Crystal structural homologues were manually docked into the low-resolution EM maps and the fit refined using the ‘Fit in Map’ function. Fits were checked by first making a low resolution map of the crystal structure (‘molmap’) and then applying ‘measure correlation’. For MW estimates, protein mass (in Da) was calculated for the estimated lower and upper contour level limits using the following calculation: 825 * V, where V is the volume (in nm^3^) of the model density at the specific contour level. See Erickson (2009) (44) for details on the calculation.

### Data availability

Data supporting the findings of this manuscript are available from the corresponding author upon reasonable request. The maps have been deposited in the Electron Microscopy Data Bank (http://www.ebi.ac.uk/pdbe/emdb/) (45): *Mycobacterium smegmatis* encapsulin: EMD-4175; *Mycobacterium smegmatis* DyP type peroxidase: EMD-10004; *Mycobacterium smegmatis* with DyP type peroxidase bound on 3-fold axis: EMD-10008; *Mycobacterium smegmatis* glutamine synthetase: EMD-4186; *Mycobacterium smegmatis* bacterioferritin: EMD-10005. Protein sequences are available from Mycobrowser (https://mycobrowser.epfl.ch/) (16): *M smegmatis* glutamine synthetase I (MSMEG_3828), bacterioferritin (MSMEG_3564), encapsulin (MSMEG_5830), and dye-decolourising type peroxidase (MSMEG_5829).

## Supplementary Methods

### Phylogenetic Analysis

Alignments of protein sequences were produced in UCSF-Chimera (43) and exported to MEGA6 (49) for phylogenetic analysis. Alignment of DNA sequences was completed in MEGA6 using MUSCLE (50) with default parameters. For protein sequences, a neighbour joining-tree was produced using p-distance to model amino acid substitution; the rate of substitution was assumed to be uniform and the pattern among lineages homogenous; gaps or missing data were deleted in the analysis. For DNA sequences, a minimal evolution tree was constructed using p-distance to model nucleotide substitutions; only transitions were included while the rate of substitution was assumed to be uniform and homogenous across lineages; gaps or missing data were deleted from the analysis. Trees were bootstrapped using 1000 replicates.

## Acknowledgments

We acknowledge Mohammed Jaffer and Brandon Weber for technical assistance and the Blackburn Group, University of Cape Town and the Yale MS & Proteomics Resource, Yale School of Medicine, New Haven, USA for mass-spectrometry measurements. Thanks to A. Mulelu for helpful discussions. This work was funded by a Research Career Advancement Fellowship from the National Research Foundation of South Africa (92556) and GCRF-START (UK Science and Technology Facilities Council grant ref. ST/R002754/1).

## Author Contributions

AMK and JDW conceived the project; AMK performed the experiments and wrote the manuscript; JDW supervised the project.

## Supplementary Figures

**Figure S1.**
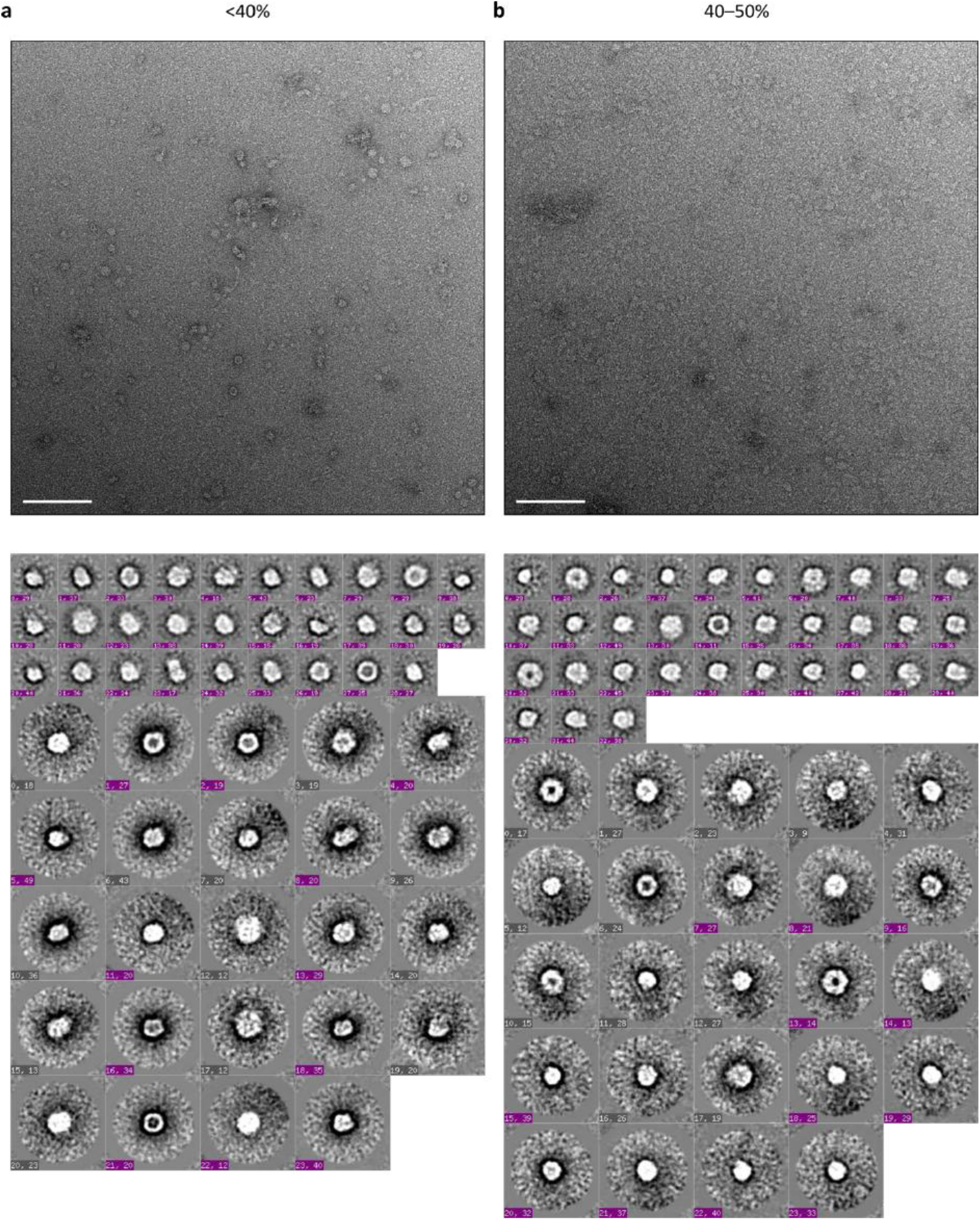
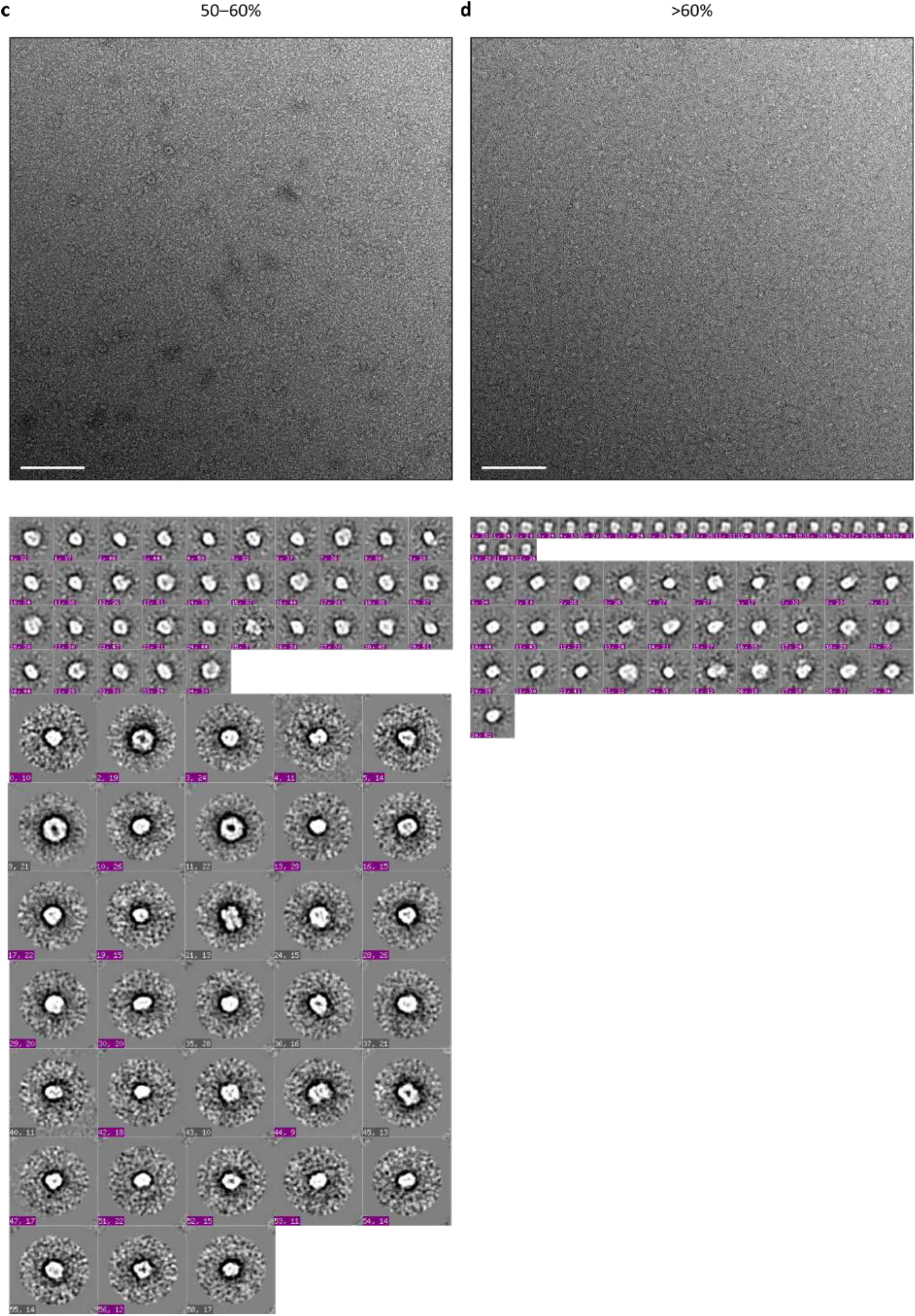
(previous page). Diversity of protein complexes in M smegmatis. Cell lysate was fractionated by a) <40%, b) 40-50%, c) 50-60%, and d) >60% ammonium sulphate cuts (top row). Particles were picked and assigned to class averages using multivariate statistics through the processing pipeline Appion (bottom row) (37) Images were taken at x50,000 magnification at a defocus of 2.00 μm using an F20 Tecnai TEM. Scale bars (white) show 100 nm.

**Figure S2.**
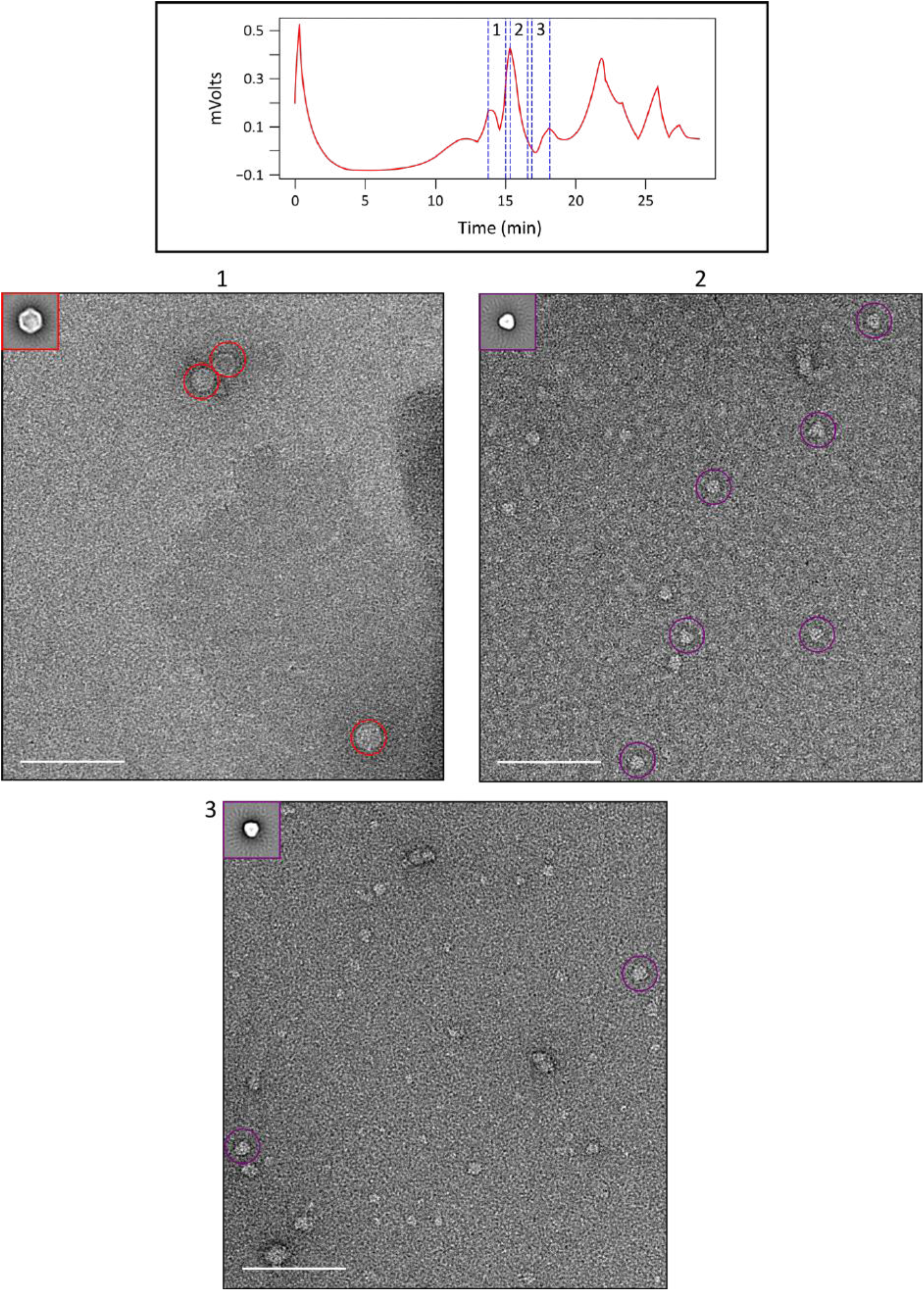
Fractionation using gel filtration after anion exchange. Three fractions (1–3) were examined from gel filtration of peak 1 (fractions #15—19) from anion exchange. Fraction 1 (#44–48) contained the presence of Encapsulin (red, circled), while fractions 2 (#49–53) and 3 (#54–58) contained a triangle-shaped average protein complex (purple, circled). Reconstruction of this complex was not pursued due to the presence of preferred orientation. The average for each particle is given in the top left-hand corner. The white scale bar shows 100 nm. Negative stain electron micrographs were taken at a magnification of x50,000 with a defocus of 2.00 μm on an F20 Tecnai TEM.

**Figure S3.**
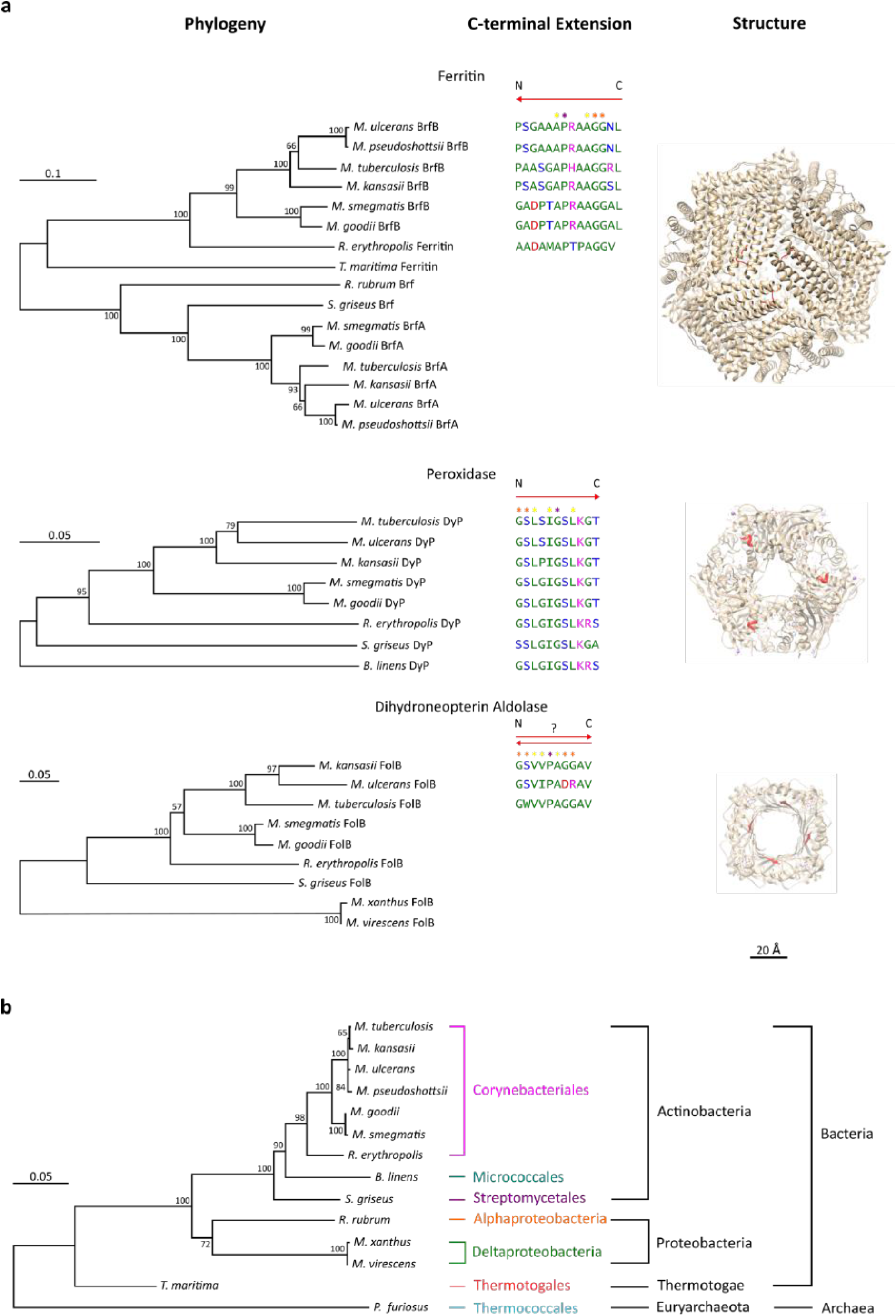
(previous page). Cargo proteins of encapsulin. **a)** The phylogeny, C-terminal extension, and structure are given for the three known cargo proteins of *Mtb.* Binding of the cargo protein to the inside of encapsulin is determined by the C-terminal extension, which is dominated by non-polar amino acids (green) with interspersed with mostly conserved polar (blue), positively charged (pink), or negatively charged (red) amino acids. The direction of binding is determined by two N- or C-terminal residues (orange star) while a central residue (purple star) separating two hydrophobic residues (yellow star) aids in positioning (21). The direction of binding for the 7,8-dihydroneopterin aldolase cargo is ambiguous. Binding of the C-terminal extension (red) is hypothesised to occur along either the 3-fold or 4-fold axis of the cargo protein. Ferritin cargo protein may also bind along its 2-fold axis (not shown). Note that the C-terminal extension is only visible for *Mtb* ferritin (pdb code 3uno (20)) and was not built into the crystal structure of peroxidase (pdb code 2gvk (46)) and was cleaved from 7,8-dihydroneopterin aldolase (pdb code 1nbu (48)). Also note that the peroxidase shown is from the closest structural homolog, *Bacteroides thetaiotaomicron.* **b)** Phylogenetic relationship between organisms that harbour known and putative encapsulin and cargo proteins, based on 16S rRNA gene sequence. While the peroxidase cargo is found in the Actinobacteria phylum, the ferritin cargo is restricted to the Corneybacteriales order, and the 7,8-dihydroneopterin aldolase cargo is specific to slow-growing Mycobacteria. For the phylogenetic trees, scale bars show amino acid or nucleotide substitutions.

**Figure S4.**
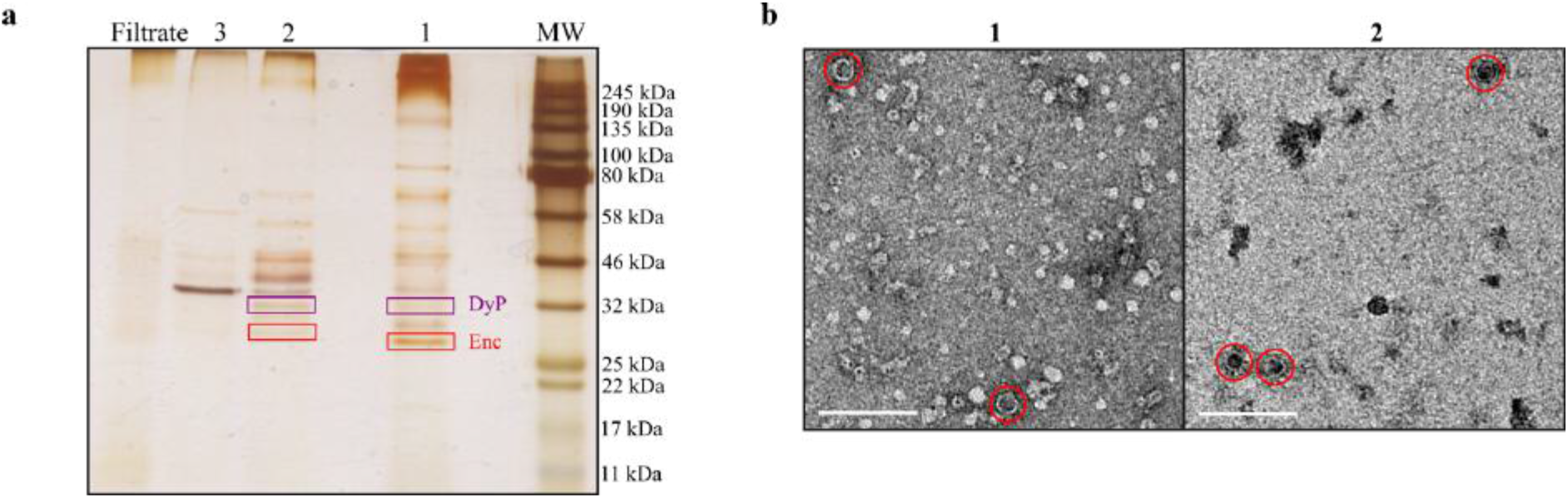
Analysis of cell culture filtrate. **(a)** Cell culture filtrate was separated by anion exchange which yielded three fractions (1–3). A silver-stained 8–15% gradient SDS-PAGE gel showed that fractions 1 and 2 contained the predominant amount of encapsulin. A mass corresponding to that of bacterioferritin B (20 kDa) was not visible. Since bacterioferritin B is present in 24 copies while DyP has 6 copies per biological unit, based on their respective symmetries, and DyP is clearly present in the gel, this suggests that DyP is encapsulated at a greater rate than bacterioferritin B. (**b**) We examined the fractions on an EM micrograph which revealed the presence of encapsulin (red circle). Images were taken at a magnification of x53,000 on the T20 Technai TEM. Scale bars show 100 nm.

